# Effective gene expression prediction from sequence by integrating long-range interactions

**DOI:** 10.1101/2021.04.07.438649

**Authors:** Žiga Avsec, Vikram Agarwal, Daniel Visentin, Joseph R. Ledsam, Agnieszka Grabska-Barwinska, Kyle R. Taylor, Yannis Assael, John Jumper, Pushmeet Kohli, David R. Kelley

## Abstract

The next phase of genome biology research requires understanding how DNA sequence encodes phenotypes, from the molecular to organismal levels. How noncoding DNA determines gene expression in different cell types is a major unsolved problem, and critical downstream applications in human genetics depend on improved solutions. Here, we report substantially improved gene expression prediction accuracy from DNA sequence through the use of a new deep learning architecture called Enformer that is able to integrate long-range interactions (up to 100 kb away) in the genome. This improvement yielded more accurate variant effect predictions on gene expression for both natural genetic variants and saturation mutagenesis measured by massively parallel reporter assays. Notably, Enformer outperformed the best team on the critical assessment of genome interpretation (CAGI5) challenge for noncoding variant interpretation with no additional training. Furthermore, Enformer learned to predict promoter-enhancer interactions directly from DNA sequence competitively with methods that take direct experimental data as input. We expect that these advances will enable more effective fine-mapping of growing human disease associations to cell-type-specific gene regulatory mechanisms and provide a framework to interpret *cis*-regulatory evolution. To foster these downstream applications, we have made the pre-trained Enformer model openly available, and provide pre-computed effect predictions for all common variants in the 1000 Genomes dataset.

**One-sentence summary:** Improved noncoding variant effect prediction and candidate enhancer prioritization from a more accurate sequence to expression model driven by extended long-range interaction modelling.

## Introduction

Models that predict gene expression and chromatin states from DNA sequence hold the promise to better understand transcriptional regulation and how it is affected by the many noncoding genetic variants associated with human diseases and traits. These models complement population-based association studies, which are limited to common variants and struggle to disentangle causality from association due to linkage disequilibrium (LD). Additionally, experimental validation of human genetic variants is laborious and limited to cell types or tissues that can be recapitulated in the lab, making it intractable to test all variants of interest in the relevant biological contexts. Although sequence-based computational models can in principle overcome these challenges, their accuracy is still limited^1–4^, making expression prediction from sequence a critical unsolved problem.

Deep convolutional neural networks (CNNs) achieve the current state of the art at predicting gene expression from DNA sequence for the human and mouse genomes^1–4^. However, to make predictions, these models are only able to consider sequence elements up to 20 kb away from the transcription start site (TSS) because the locality of convolutions limits information flow in the network between distal elements. Many well-studied regulatory elements, including enhancers, repressors, and insulators, can influence gene expression from far greater than 20 kb away^5^. Thus, increasing information flow between distal elements is a promising path to increase predictive accuracy.

## Results

### Enformer improves gene expression prediction in held-out genes

We developed a new model architecture named Enformer (a portmanteau of enhancer and transformer) to predict gene expression and chromatin states in human and mouse from DNA sequence (**Fig. 1a, Extended Data Fig. 1**). Transformers are a class of deep learning models that have achieved significant breakthroughs in natural language processing (NLP)^6,7^. They consist of attention layers that transform each position in the input sequence by computing a weighted sum across the representations of all other positions in the sequence. Attention weight between any two positions depends on the embeddings of their current representation vectors and the distance between them. This allows the model, for example, to refine the prediction at a transcription start site (TSS) by gathering information from all relevant regions, such as enhancers regulating the gene. Since each position directly attends to all other positions in the sequence, they allow for a much better information flow between distal elements. By contrast, convolutional layers require many successive layers to reach distal elements due to their local receptive field. Using transformer layers allowed us to nearly double the receptive field, reaching distal regulatory elements up to 100 kb away while still being able to effectively integrate their information. By contrast, previous state-of-the-art models Basenji2 or ExPecto only reach elements up to 20 kb away (**Extended Data Fig. 1**). This increase in the receptive field is important because it greatly expands the number of relevant enhancers seen by the model from 47% (<20kb) to 84% (<100 kb) as estimated from the proportions of high-confidence enhancer-gene pairs^8^.

**Fig. 1:**
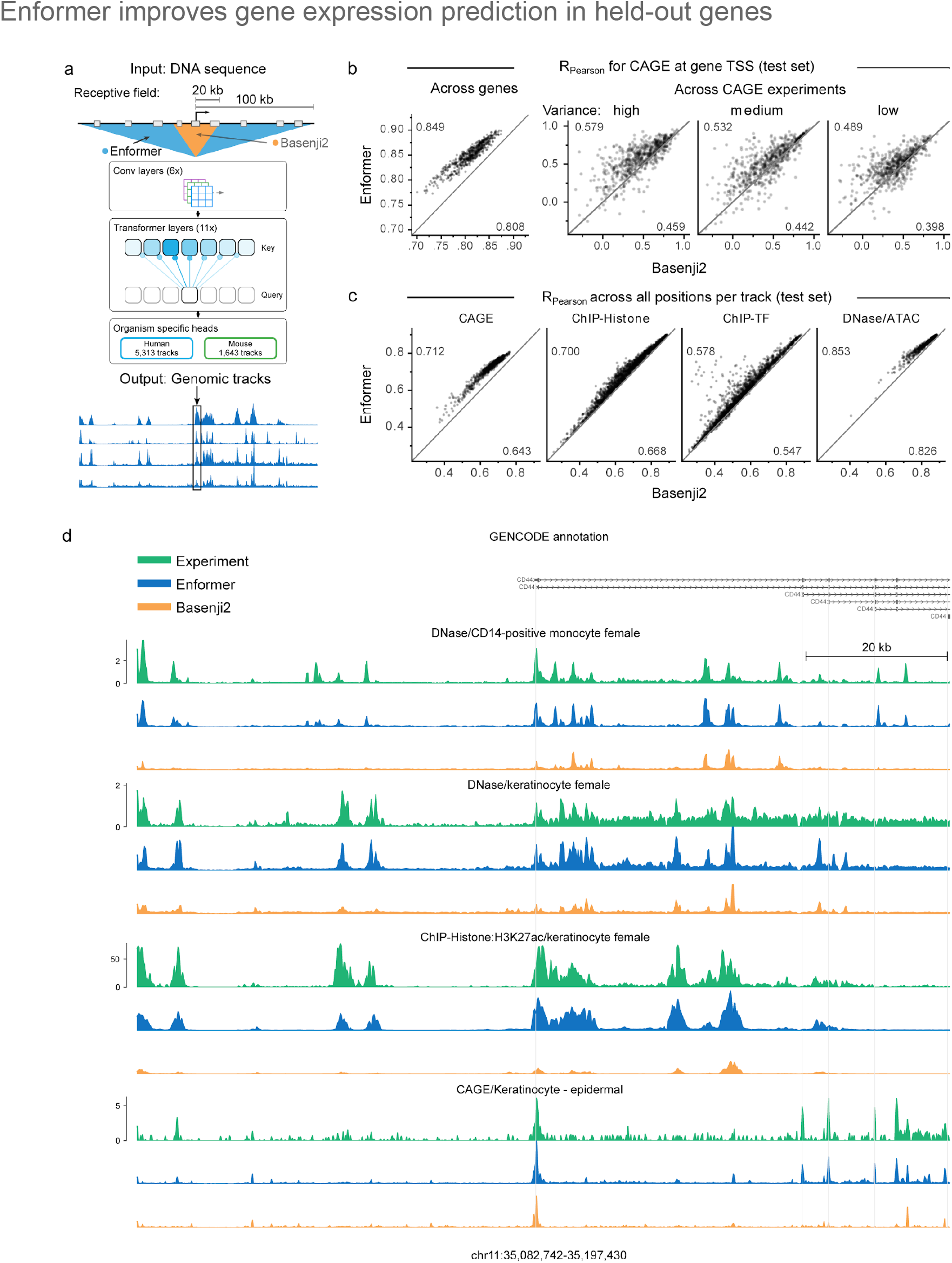
Enformer improves gene expression prediction in held-out genes by using a larger receptive field. **a)** Enformer is trained to predict human and mouse genomic tracks at 128 bp resolution from 200 kb of input DNA sequence. By using transformer modules instead of dilated convolutions, it achieves a five times larger receptive field able to detect sequence elements 100 kb away compared to 20 kb for Basenji2^2^ or ExPecto^1^ (**Extended Data Fig. 1**). **b)** Enformer outperforms Basenji2 in gene expression prediction from sequence both across genes and across CAGE experiments for protein-coding genes. Test set performance was measured by Pearson correlation of CAGE gene expression (log(1+x) transformed) computed across genes for each CAGE experiment (left) or across CAGE experiments for each test gene stratified by the observed expression variance across experiments (right). Average performance for each model is shown in the corners. Bootstrapped standard deviation of these estimates is 0.004 for ‘Across genes’. Gene expression values were obtained by summing up the observed or predicted CAGE read counts at all unique TSS locations of the gene. Values for each CAGE experiment were standardized to have zero mean and variance of 1 across genes. **c)** Enformer consistently outperforms Basenji2 across all 4 assay types (columns) as measured by Pearson correlation computed across all 128 bp binned genomic positions in the human test set for 5,313 predicted tracks (points). Both models were trained and evaluated on the same dataset. **d)** Representative example of observed and predicted genomic tracks (log_10_ scale) at CD44 gene locus located in the test set region with high disagreement between Enformer and Basenji2 predictions. For each experiment, all three tracks share the same y-axis.

Enformer substantially outperformed the previous best model Basenji2 for predicting RNA expression as measured by Cap Analysis Gene Expression^9^ (CAGE) at the TSS of human protein-coding genes, with the mean correlation increasing from 0.81 to 0.85 (**Fig. 1b**, left). This performance increase is twice as large as the performance increase between Basenji1^3^ and Basenji2^2^ and closes a third of the gap to the experimental-level accuracy estimated at 0.94 (**Extended Data Fig. 2a**). Gene expression predictions also better captured tissue or cell-type specificity (**Fig. 1b**, right). The performance improvement was consistent across all four types of genome-wide tracks including CAGE measuring transcriptional activity, histone modifications, TF binding, and DNA accessibility in various cell types and tissues for held-out chromosomes (**Fig. 1c**). The performance improvement was largest for CAGE, possibly because tissue-specific gene expression strongly depends on distal elements^10^. The improvement in prediction accuracy was also qualitatively evident when visualizing observed and predicted tracks of the genome (**Fig. 1d**). These results confirm that the Enformer architecture is well suited to predict both the broad range of epigenetic marks as well as gene expression from DNA sequence.

To pinpoint the benefit of attention layers compared to the dilated convolutions used in Basenji2, we replaced attention layers with dilated convolutions and tuned the learning rate for optimal performance. Attention layers outperformed dilated convolutions across all model sizes, number of layers, and number of training data points (**Extended Data Fig. 2b**). The larger receptive field was indeed key, because we observed a large performance drop when restricting the receptive field of Enformer to that of Basenji2 by replacing global attention layers with local ones (**Extended Data Fig. 2c**). We note that increasing the number of parameters improved model performance, consistent with recent advances in NLP^7^. Enformer uses custom relative positional basis functions in the transformer layers to more easily distinguish between proximal and distal enhancers, and to distinguish positions upstream and downstream of the TSS. Both properties provided a noticeable performance improvement over the typically used relative basis functions and absolute positional encodings in the NLP literature (**Extended Data Fig. 3a,b**). Overall, these results confirm that attention layers are better suited than dilated convolutions for gene expression prediction.

### Enformer attends to cell-type-specific enhancers

To better understand what sequence elements Enformer is utilizing when making predictions, we computed two different gene expression contribution scores―input gradients (gradient*input)^12^ and attention weights (Methods)―for several genes with most CRISPRi-validated enhancers^8,11^. Contribution scores highlight the input sequences most predictive for the expression of a particular gene in a given cell type^13,14^. We inspected the contribution scores of several genes and observed that they correlated with H3K27ac and highlighted not only local promoter regions, but also distal enhancers more than 20 kb away (Fig. 2a, Extended Data Fig. 4, 5). By contrast, the contribution scores of Basenji2 were zero for sequences beyond 20 kb from the TSS due to the limited receptive field, thereby missing several enhancers. This example suggests that Enformer is indeed looking at biologically relevant regions such as enhancers beyond 20 kb when making predictions, and that gene expression contribution scores could be used to prioritize relevant enhancers.

**Fig. 2:**
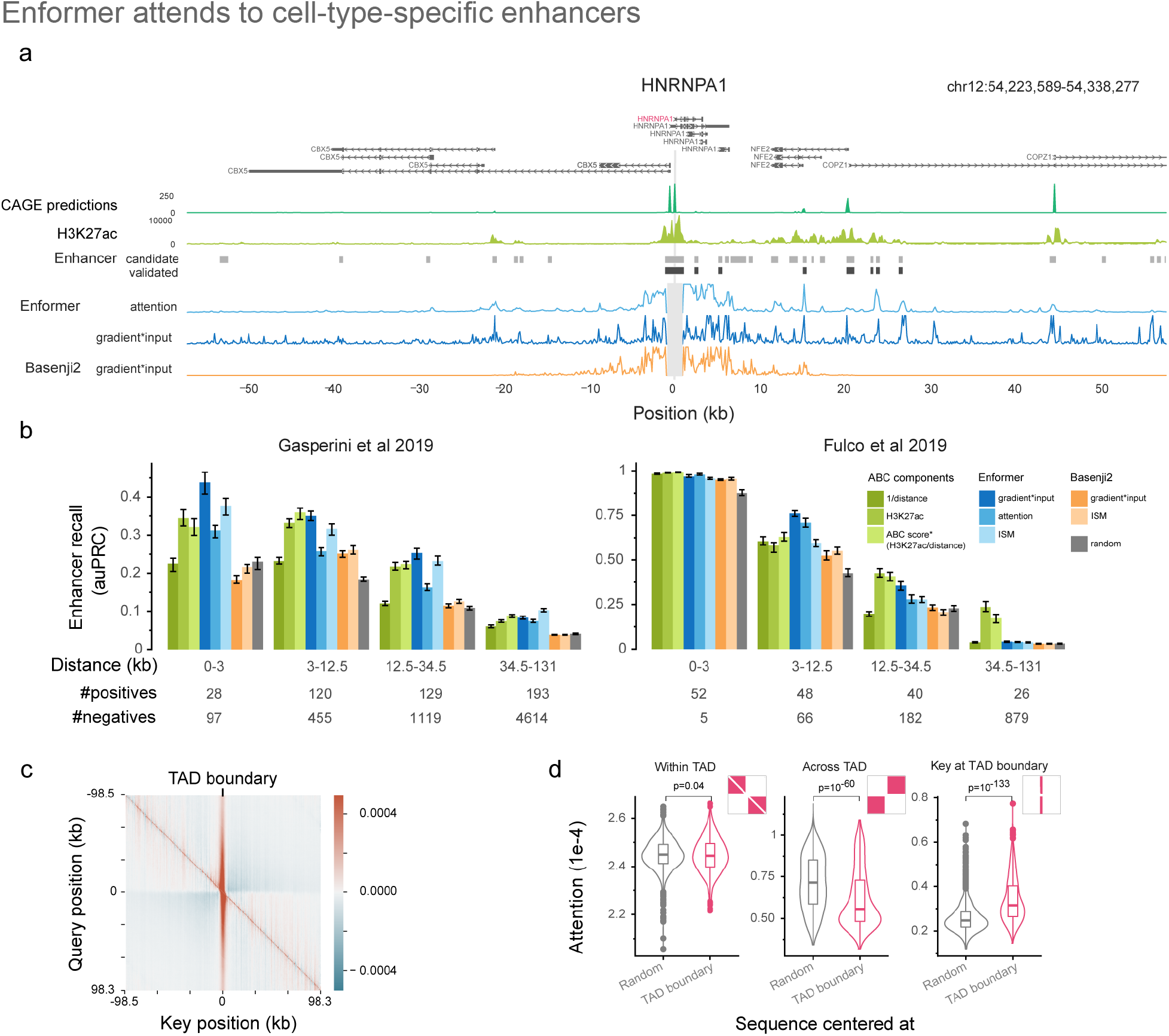
Enformer attends to cell-type-specific enhancers when making predictions, enabling enhancer prioritization. **a)** Example gene HNRNPA1 locus showing from top to bottom: gene annotations; predicted CAGE expression in K562 (chronic myeloid leukemia cell line); measured H3K27ac (modification associated with active enhancers) in K562; candidate (light gray) and CRISPRi-validated enhancers (dark gray) exhibiting significant HNRNPA1 expression changes from Fulco et al 2019^11^; Enformer attention weight averaged across all layers and heads for a query placed at the main TSS of HNRNPA1 gene (position 0); and gradient*input^12^ contribution scores computed w.r.t. the K562 CAGE track at the main TSS position for Enformer and Basenji2. **b)** Enhancer–gene pair classification performance (CRISPRi-validated vs non-validated candidate enhancers), stratified by relative distance, as measured by area under precision-recall curve (auPRC) on two CRISPRi datasets^8,11^ for different methods, models, and contribution scores (Methods). ABC score* (H3K27ac/distance) denotes the approximate version of the ABC score^11^ lacking DNase and Hi-C data while exhibiting similar or better performance (**Extended Data Fig. 6a**). Error bars show the 25th and 75th percentile of the auPRC distribution obtained by sampling 80% of enhancer–gene pairs (100 times without replacement). The number of positive/negative enhancer–gene pairs are shown at the bottom. The auPRC metric is sensitive to class imbalance, hence the absolute auPRC numbers between the two datasets or different distance windows are not directly comparable. For example, Gasperini et al. 2019^8^ is more imbalanced: 1:10 vs 1:4 in Fulco et al. 2019^11^. **c)** Average attention matrix difference of Enformer between 1,500 sequences centered at a topologically associating domain (TAD) boundary (position=0) and 1,500 sequences from the human validation set without any particular centering. Attention matrices were averaged across all layers, heads, and sequences. Red and blue denote regions where attention is higher and lower, respectively, for TAD-boundary sequences compared to randomly centered sequences. Red stripe in the center at key=0 means that the model is attending to the TAD boundary more strongly than by chance. Blue regions in off-diagonal quadrants means that model is attending less across the TAD-boundary, consistent with the biological observation of reduced inter-TAD interactions. **d)** Attention is significantly lower across TAD boundaries (center), significantly higher at TAD boundaries (right), and shows no significant difference within them (left). Shown is the distribution of average attention values across all sequences in different parts of the attention matrix (visualized as a mask in the top right corner). P-values were computed with the two-sided Mann–Whitney U test. The box plots mark the median, upper and lower quartiles and 1.5x interquartile range (whiskers); outliers are shown as points.

Linking candidate enhancers identified via biochemical annotations^15^ to target genes is an important and unsolved problem^5^. Computational models have historically produced low accuracy due to the combination of noisy labels and class imbalance. To systematically evaluate the ability of contribution scores to pinpoint relevant enhancers for a particular gene, we compared several contribution scores across all tested enhancer–gene pairs in two large-scale CRISPRi studies performed on the K562 cell line^8,11^. In these experiments, CRISPRi was used to suppress the activity of more than 10,000 candidate enhancers and measure their effect on gene expression.

Enformer contribution scores prioritized validated enhancer–gene pairs with higher accuracy than Basenji2 contribution scores or random scores across almost all relative distances and different types of contribution scores (**Fig. 2b**, Enformer vs Basenji vs Random). The performance of Enformer was comparable to, and in some cases even better than, the ABC score^11^, a state-of-the-art method recently proposed specifically for enhancer prioritization. This is remarkable because the ABC score relies on experimental data such as a HiC-based interaction frequency and H3K27ac as input (**Fig. 2b**: blue vs green, **Extended Data Fig. 6a**), whereas Enformer only uses DNA sequence as input and was never trained to explicitly locate enhancers. This allows Enformer to also be used for arbitrary sequence variations lacking experimental data. High prioritization performance for Enformer was only obtained for contribution scores that were cell-type-specific (**Extended Data Fig. 6c**). Thus, Enformer contribution scores are an effective strategy to prioritize candidate enhancers in cell types used for model training.

Next, we asked whether the model has learned about another important class of regulatory elements: insulator elements, which separate two topologically associating domains (TADs) and minimize enhancer–promoter crosstalk between the two. We inspected the attention matrices (which were more efficient to compute relative to input gradients due to many output targets) of sequences centered at TAD boundaries, and compared them with attention from sequences with no particular alignment. From the perspective of the query position, Enformer paid more attention to TAD boundaries compared to random positions (vertical red stripe, **Fig. 2c**) and less attention to regions on the opposite side of the boundary (off-diagonal blue blocks, **Fig. 2c**). Both of these two patterns were statistically significant across 1,500 tested sequences (**Fig. 2d**, “Across TAD” and “Key at Insulator”). This means that the model has not only learned about the role of tissue-specific enhancers and promoters, but also about insulator elements and their role in inhibiting information flow between two genomic compartments.

### Enformer improves variant effect prediction on eQTL data

A central goal of this research is to predict the influence of genetic variants on cell-type-specific gene expression, in order to inform fine-mapping of the many thousands of noncoding associations with phenotypes of interest from genome-wide association studies (GWAS). Computational models that predict regulatory activity from DNA sequence can process distinct alleles and compare predictions to score genetic variants^3,18–20^. A successful model would be able to produce the results of a gene expression quantitative trait loci (eQTL) study without having to measure hundreds to thousands of individual gene expression profiles. Thus, we studied eQTLs discovered by the GTEx project across dozens of human tissues to validate model predictions^21^. The primary challenge of such validation is the influence of co-occurrences between variants (i.e. linkage disequilibrium) in the profiled population, which transfers the causal eQTL’s effect to co-occurring variants. Signed linkage disequilibrium profile (SLDP) regression is a technique developed to measure the genome-wide statistical concordance between signed variant annotations (such as our model predictions) and GWAS summary statistics (such as GTEx eQTLs) while accounting for linkage disequilibrium (Methods)^22^. For 379 of 648 (59.4%) CAGE datasets, the maximum SLDP Z-score across GTEx tissues (representing the most likely closest sample match) increased for Enformer predictions relative to Basenji2. Enformer maximum Z-scores increased by greater than one standard deviation for 228 CAGE datasets, relative to 46 decreased by one. The maximum Z-score increased on average from 6.3 to 6.9 (**Fig. 3a**). Note that we do not expect increased SLDP Z-scores for CAGE samples without a relevant GTEx tissue match. We observed a qualitative improvement in the tissue similarity of the top ranked CAGE sample for GTEx tissues, exemplified by increased SLDP Z-scores for muscle samples to GTEx skeletal muscle and adipose samples for GTEx subcutaneous adipose tissue (**Fig. 3b,c**). Thus, Enformer predictions for noncoding variant activity appear to improve primarily for samples with similar cell type composition, in line with our observations of improved tissue and cell-type specificity for held-out sequences.

**Fig. 3:**
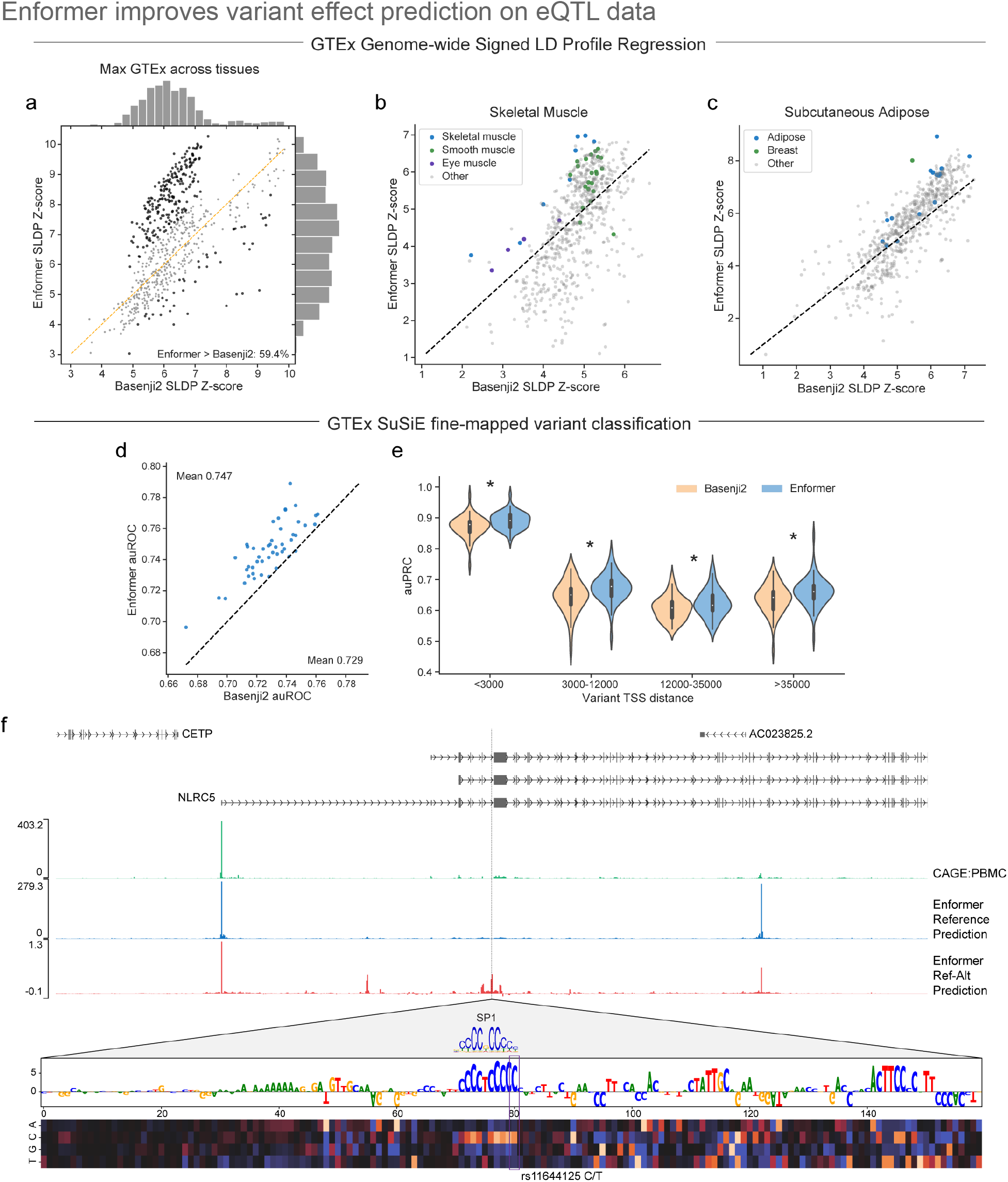
Enformer improves variant effect prediction on eQTL data as measured by SLDP regression and fine-mapped variant classification. **a)** We computed genome-wide statistical concordance between variant effect predictions for individual CAGE datasets and GTEx eQTL summary statistics using SLDP across all variants in the 1000 genomes dataset. Taking the max tissue for each sample, Enformer predictions achieved greater Z-scores for 59.4% of samples, and 228 are greater by more than one standard deviation (versus 46 for Basenji2). **b, c)** Studying SLDP in skeletal muscle (b) and subcutaneous adipose (c) GTEx tissues indicated that biologically relevant CAGE datasets (shown in blue) improve between Basenji2 and Enformer. **d)** We trained random forest classifiers to discriminate between fine-mapped GTEx eQTLs and matched negative variants in each of 48 tissues (Methods). Features derived from Enformer enabled more accurate classifiers than Basenji2 features for 47/48 tissues. **e)** We computed auPRC for variants in four roughly equally sized TSS distance bins. Violin plots represent measures for the 48 tissues. Enformer improved accuracy at all distances (Wilcoxon p-values < 1e-4). **f)** Enformer prediction for rs11644125 improved relative to Basenji (data not shown) by better capturing its influence on an NLRC5 TSS ∼35 kb upstream. rs11644125 is associated with monocyte and lymphocyte counts in the UK BioBank and fine-mapped to >0.99 causal probability^16^. In silico mutagenesis of the region surrounding rs11644125 revealed an affected SP1 transcription factor motif^17^.

Although linkage disequilibrium generally results in GTEx eQTL associations that can only be attributed to a set of frequently co-occurring variants, the latest GTEx release includes many thousands of associations in loci with simple linkage patterns, which have been fine-mapped to a single high probability causal variant^23^. To assess the utility of Enformer predictions for identifying of causal variants, we defined a classification task for each tissue to discriminate likely causal variants (causal probability>0.9, as determined by the population-based fine-mapping model SuSiE^24^) from likely spurious eQTLs (causal probability<0.01), which were matched for the eGene when possible (Methods). We represented each variant by its prediction difference vector (i.e. evaluating the reference minus alternative allele, summed across the sequence) for all 5,313 human data sets, and trained random forest classifiers. Enformer predictions enabled a more accurate classifier for 47/48 GTEx tissues (**Fig. 3d**), increasing the mean auROC from 0.729 to 0.747. This improvement was consistent across all distances from the TSS (**Fig. 3e**), suggesting that the model not only better represents variants likely overlapping long-range enhancers (enabled by the larger receptive field), but also more effectively parses promoters and short-range enhancers. The Enformer model was also more accurate at predicting the direction of expression change of these fine-mapped eQTLs than Basenji2 (**Extended Data Fig. 7**).

One example variant where the Enformer eQTL probability prediction increased relative to Basenji2 is rs11644125, which lies within an intron ∼35 kb downstream of the TSS of NLRC5, a protein involved in viral immunity and the cytokine response (**Fig. 3f**). The variant has been statistically fine-mapped as likely to cause changes in monocyte and lymphocyte blood cell counts^16^. According to GTEx, the minor allele T decreases gene expression of NLRC5 in whole blood relative to the major allele C. Enformer correctly predicts reduced NLRC5 expression from the upstream TSS in many relevant CAGE samples, including PBMCs. Using *in silico* mutagenesis of the local region (Methods), we observed that the variant rs11644125, modulates the known motif of the transcription factor SP1^16^. Enformer predictions suggest perturbed SP1 binding in hematopoietic cells that alters NLRC5 expression as a mechanism for these traits.

### Enformer outperforms winning solution for critical assessment of genome interpretation (CAGI5) challenge

Finally, we evaluated Enformer’s performance on a second, independent variant effect prediction task using a dataset in which massively parallel reporter assays (MPRAs) directly measured the functional effect of genetic variants through saturation mutagenesis of several enhancers and promoters in a variety of cell types^25^. We used the same training and test sets as the CAGI5 competition^26^, enabling us to directly benchmark Enformer’s performance relative to those of submissions from other groups. For each variant, we evaluated its effect as the predicted difference between the reference and alternative allele, retrieving 5,313 features. Next, we compared two approaches: i) we used these features to train a lasso regression model on the provided training set for each gene, and ii) we pre-selected a subset of features corresponding to cell-type matched and cell-type agnostic predictions of changes in CAGE and DNase, and generated a summary statistic of the features (i.e. without additional training).

Evaluating these two approaches on each gene’s test set revealed that lasso regression with Enformer predictions as features had the best average correlation across all loci, among seven alternative submissions from the competition (**Fig. 4a**). Moreover, using the Enformer predictions directly as scores, without training, performed comparably to the lasso-trained model and also outperformed the other submissions. This includes the sequence-based predictor deltaSVM^27^, which was trained on independent DNase and H3K4me1 data derived from matched cell types^25^. The lasso-trained Enformer exceeded the performance of Group 3, the winning team from CAGI5 (p<0.01, paired, one-sided Mann–Whitney U test, **Fig. 4b**). Visualization of the predictions that required no additional training revealed that Enformer faithfully captured the effects of two out of four transcription factor binding sites for the LDLR locus (**Fig. 4c**). Enformer highlighted an additional binding site that had lower effect sizes, but still showed a significant difference. By contrast, deltaSVM successfully predicted only one binding site but missed the other three, overall exhibiting 50% reduced Pearson and Spearman correlations to the measured effects relative to Enformer. For this locus, cell-type matched predictions mirrored cell-type agnostic predictions, indicating that the binding sites which were detected likely corresponded to general transcription factors present in most cell types.

**Fig. 4:**
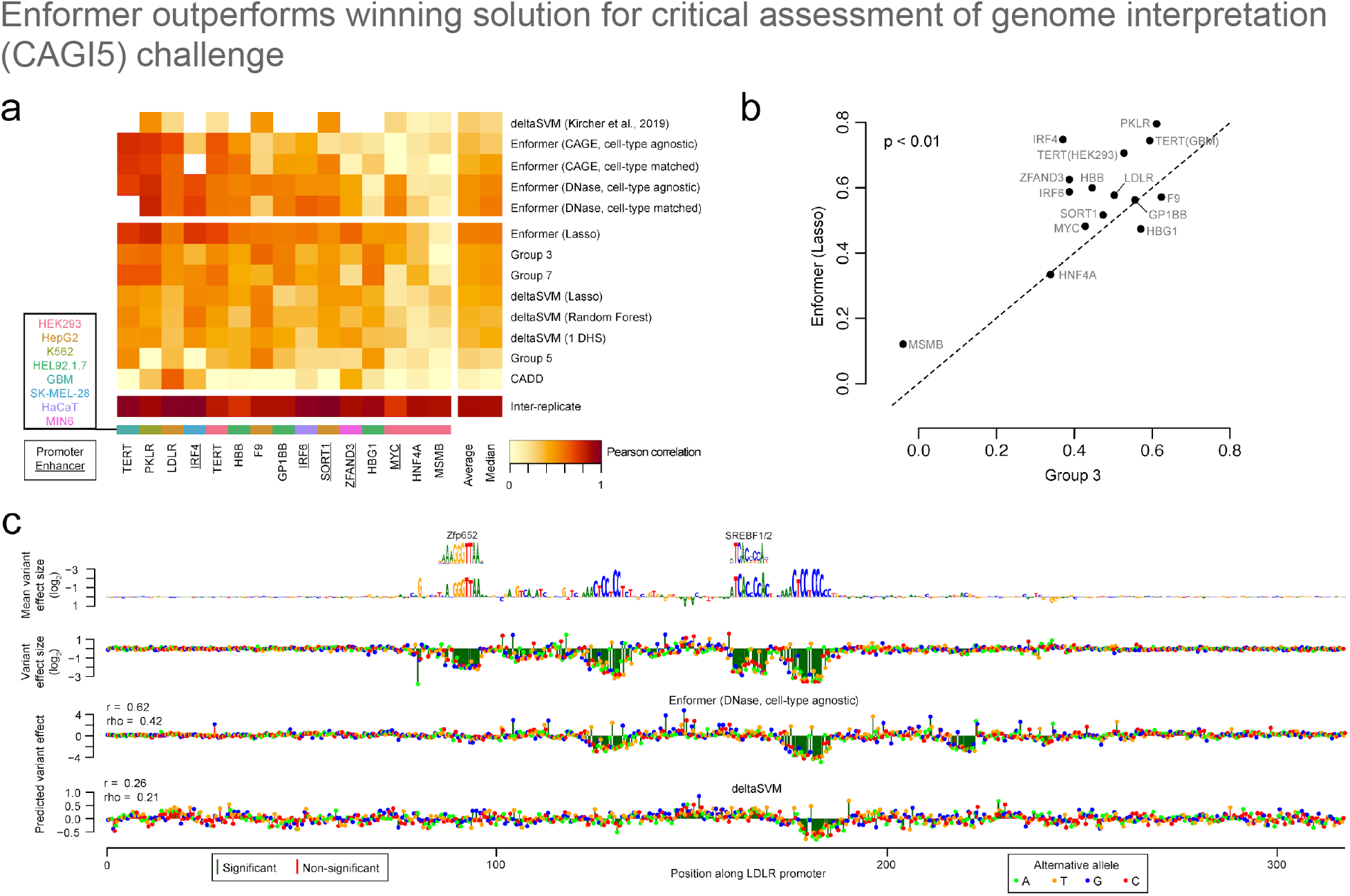
Enformer outperforms winning solution for critical assessment of genome interpretation (CAGI5) challenge. **a)** Correlation of variant effect predictions with experimental values, as measured by saturation mutagenesis MPRAs^25^, on test sets for 15 loci curated for the CAGI5 competition^26^. Shown above the horizontal break is the performance of 5 methods that required no additional fine-tuning on each locus; shown below it is that of 8 methods that were additionally trained on the CAGI5 training sets. **b)** Pearson correlations of each locus for predictions derived from the Enformer vs the winning team of the CAGI5 competition. Enformer shows a significant performance improvement (p<0.01, one-sided Mann–Whitney U test). **c)** Example saturation mutagenesis data from the LDLR promoter locus. Shown in the top row is the reference sequence scaled to the mean effect size among all alternative mutations, with measured effect sizes of individual variants in the second row. Two of the four significant elements match known motifs^17^, and the two unknown motifs partially resemble the SP1 binding motif. Shown in the bottom two rows are the predictions on the full dataset using methods from (a) that required no additional fine-tuning.

## Discussion

A long-standing problem in regulatory genomics is that of predicting gene expression purely from DNA sequence. With a novel transformer architecture, we have made a significant improvement by greatly expanding the receptive field and increasing the information flow between distal elements. In this way, the model can better capture biological phenomena such as enhancers regulating promoters despite large DNA sequence distance between the two. This led to a substantial performance increase in tissue and cell-type-specific gene expression prediction correlation from 0.81 to 0.85, one third of the way towards the experimental-level accuracy of 0.94 estimated from replicates.

This improvement in predictive accuracy translated to improved models for two key problems of biological relevance: enhancer–promoter prediction and noncoding variant effect prediction. We observed that the model pays attention to enhancers and considers insulators when making gene expression predictions, suggesting that it has learned canonical distal regulation patterns. Using the Enformer model, we can more accurately predict whether a natural variant or CRISPR-perturbed enhancer will cause a significant expression change compared to previous approaches. By relying solely on DNA sequence as input, Enformer has several advantages over alternative variant effect prediction methods: i) unlike most methods^25^, it is capable of signed prediction of activating or repressive mutations, ii) by not relying explicitly on nucleotide conservation statistics as the majority of tools do^25^, its predictions are not limited to conserved enhancers, which comprise a small proportion of all enhancers^28^, and iii) it can make predictions for arbitrary sequences, which enables the synthetic design of enhancers that are optimized to exhibit cell-type specificity^29^. Altogether, these advances and advantages open exciting avenues to study the expanding catalogs of genetic variants linked to disease and enhancer biology in development and evolution.

Several paths to further improve model accuracy appear promising. Machine learning success depends on the training data, so increasing the resolution and quality of the target tracks^14^, and curating data from additional organisms^2^, would likely boost performance. Recent work demonstrated that the highly structured 3D DNA contacts, which greatly influence long-range gene regulation, are predictable from the underlying DNA sequence^30,31^. Artful combination of these models with our own could improve Enformer’s modeling of insulators and distal regulation. A limitation of the current approach is that we can only model and predict for cell types and assays in the training data and cannot generalize to new cell types or assays. Parallel research has begun to address this shortcoming via representation learning of cell types and assays and could make use of the Enformer architecture in the future^32,33^. The sensitivity of the model to genetic variants could be further improved by training upon the growing number of functional genomic datasets, such as those derived from CRISPR perturbations and massively parallel reporter assays. Currently, the small size of these datasets has limited their usage only to model evaluation. Finally, we anticipate that recent improvements in the computational efficiency^34^ of transformer models together with better hardware will allow us to further scale-up the models.

In the future, Enformer could be systematically applied to fine-map existing GWAS studies^23^, prioritize rare or *de novo* variants observed for rare disorders^35,36^, and impute regulatory activity across species to study *cis*-regulatory evolution^2^. To foster these downstream applications, we have made the pre-trained Enformer model openly available and have also provided a [Colab notebook](https://colab.research.google.com/github/deepmind/deepmind_research/blob/master/enformer/enformer-usage.ipynb) demonstrating its use. Furthermore, we have pre-computed effect predictions for all frequent variants in the 1000 Genomes dataset and made them openly available here. To make these predictions more accessible, we distilled the 5,313 features into 20 highly informative variant scores using PCA (Methods) to keep the released file sizes manageable (<1 GB in total for 10 M variants, instead of 100 GB) while retaining high predictive accuracy (GTEx fine-mapping classification auROC of 0.743 compared to 0.747 using all features). We hope that our model will stimulate an improved understanding of gene regulatory architecture and facilitate the development of improved diagnostic tools for diseases of genetic origin.

## Data availability

Gene annotation was obtained from https://www.gencodegenes.org/. Basenji2 training, validation, and test data was obtained from https://console.cloud.google.com/storage/browser/basenji_barnyard/data. Processed CRISPRi data for Fulco et al 2019^11^ was obtained from supplementary material and for Gasperini et al 2019^8^ from GEO accession GSE120861. H3K27ac ChIP-seq data in K562 used for analysis in Fig. 2 was obtained from https://www.encodeproject.org/ with file accession ENCFF779QTH and DNase with file accessions ENCFF413AHU and ENCFF936BDN. We downloaded GTEx summary statistics from https://gtexportal.org and SuSiE fine mapping results from Wang et al. 2020^23^ We acquired training and test sets as well as the predictive accuracies of individual competition participants from the CAGI5 competition^26^ (M. Kircher, personal communication, https://genomeinterpretation.org/content/expression-variants).

## Code availability

All components of our core algorithm, including the full model architecture and example code to train and evaluate the model are available under the open source Apache 2.0 license at the following URL: https://github.com/deepmind/deepmind-research/tree/master/enformer. In addition, layer components of the model are now available in the existing Basenji repository for biological sequence deep learning at https://github.com/calico/basenji also under the open source Apache 2.0 license.

Pre-trained enformer model is available on TF-Hub so that users can easily run it on new data: https://tfhub.dev/deepmind/enformer/1. We also plan to release it in the Kipoi model repository^37^. We provide code examples (enformer-usage.ipynb) on how to use that model to score genetic variants. Finally, we provide variant effect predictions for all frequent variants in the 1000 genomes cohort (MAF>0.5% in any population) here, with an open creative-commons CC-BY 4.0 license.

## Acknowledgements

We thank Martin Kircher for sharing saturation mutagenesis MPRA datasets and variant effect predictions with us as well as Alexander Pritzel, Alexander W. R. Nelson, Antonia Patterson, Annette Obika, Clemens Meyer, Demis Hassabis, Michelle Dunlop, Natasha Latysheva, Nazif Alic, Sara-Jane Dunn, Stig Petersen, Teresa Niccoli, Toby Sargeant and Trevor Back for their contributions and support.

## Author contributions

Ž.A., J.R.L., D.R.K., and P.K. initiated the project. Ž.A., V.A., and D.R.K. conceived of the study and designed the analyses. Ž.A. designed the model with help from D.V., J.J., and P.K.. Ž.A., D.V., K.T., and Y.A. implemented the model. Ž.A. performed model performance analyses, A.G-B. and Ž.A. performed enhancer prioritization analysis, D.R.K. performed variant effect analysis on population genetic data, and V.A. performed variant effect analyses for MPRA data. J.R.L., J.J., P.K., and D.R.K. supervised the study. Ž.A., V.A., and D.R.K. prepared the manuscript with input from all authors.

## Methods

### Model architecture

The Enformer architecture consists of three parts: i) 7 convolutional blocks with pooling, ii) 11 transformer blocks, and iii) a cropping layer followed by final pointwise convolutions branching into two organism-specific network heads (**Extended Data Fig. 1)**. Enformer takes as input one-hot-encoded DNA sequence (A=[1,0,0,0], C=[0,1,0,0], G=[0,0,1,0], T=[0,0,0,1], N=[0,0,0,0]) of length 196 kb and predicts 5,313 genomic tracks for the human genome and 1,643 tracks for the mouse genome, each of length 896 corresponding to 114,688 bp aggregated into 128 bp bins. Convolutional blocks with pooling first reduce the spatial dimensionality from 196 kb to 1,536, transformer blocks capture long-range interactions across the whole sequence (128 bp resolution), a cropping layer trims 320 positions (128 bp resolution) on each side to ensure each output position will have at least 40,960 bp of sequence context on both sides, and the output heads predict organism-specific tracks. The Enformer architecture is similar to the state-of-the-art (SOTA) model Basenji2^2^. However, the following changes helped us improve and exceed the SOTA performance: Enformer uses transformer blocks instead of dilated convolutions, attention pooling instead of max pooling, twice as many channels, and 1.5 times longer input sequence (196 kb instead of 131 kb). The detailed model architecture, including the selected hyperparameters, is shown in **Extended Data Fig. 1**.

Attention pooling summarizes a contiguous chunk of the input sequence 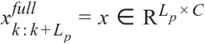 across L_p_ positions for each of the *C* channels and returns the output value **h** ∈ *R*^*C*^ as follows:

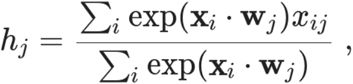

where *i* indexes sequence position weighted by the exponentiated dot product **x**_i_ ·**w**_j_ and **w** ∈ *R*^*C × C*^ is a matrix of learned weights. We apply attention pooling to contiguous chunks of the original input sequence using window size L_p_=2 and stride of 2. We initialize **w** to 2 * **1** where **1** is the identity matrix to make the operation similar to max pooling. This initialization gave slightly better performance than random initialization or initialization with zeros representing average pooling.

We use multi-head attention (MHA) layers to share information across the sequence and model long-range interactions, such as those between promoters and enhancers. Each head has a separate set of weights **w**^q^ ∈ *R*^*CxK*^, **w**^k^ ∈ *R*^*CxK*^, **w**^v^ ∈ *R*^*CxV*^ which transform the input sequence **x** ∈ *R*^*LxC*^ into queries **q**=**x w**^q^, keys **k**=**x w**^k^, and values **v**=**x w**^v^. Queries represent the current information at each position, and keys represent the information each position will be looking for to attend to. Their dot product forms the attention matrix (without relative positional encodings), which is computed as **a**=softmax(**q k**^**T**^/√K), where the entry a_ij_ represents the amount of weight query at position i puts on the key at position j. Values represent the information that each position will propagate forward to positions that attend to it. Each single attention head computes output as a weighted sum across all input positions: **av**. This allows each query position to use the information across the whole sequence. The multiple heads compute with independent parameters, and we concatenate the outputs from each head to form the final layer output followed by a linear layer to combine them. Our layers used 8 heads, value size of 192, and key/query size of 64.

To inject positional information, we augment **a** with relative positional encodings^38^ as formulated in the Transformer-XL paper^39^ without using any recurrent state. Relative positional encodings provide a parameterized baseline for how actively two positions in the sequence should influence each other during the layer’s transformation as a function of their pairwise distance. We use 3 different basis function classes, as visualized in **Extended Data Fig. 2c**:

1. 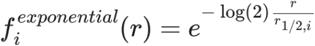, where r_1/2,i_ is placed linearly in the log-space between 3 and sequence length.
2. 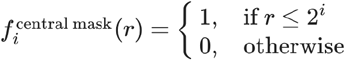
3. 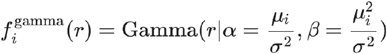, where Gamma(r|α,β) is the gamma probability distribution function. μ_i_ is placed linearly from (sequence length / number of features) to sequence length and σ=sequence length / (2 * number of features).

For each basis function, we use a symmetric f(|x|) and asymmetric sign(x) * f(|x|) version to introduce directionality. We use the same number of relative positional basis functions as the value size of MHA (192). The 192 basis functions are equally divided among the basis function classes and the symmetric vs asymmetric versions thereof. With 3 basis function classes, each basis function class provides 64 positional features (32 symmetric and 32 asymmetric).

Dropout rates of 0.01 and 0.05 were used for positional encoding features and the final attention matrix respectively in MHA. All other dropout rates are annotated in **Extended Data Fig 1a**.

## Model training and evaluation

The model was trained, evaluated, and tested on the same targets using the same Poisson negative log-likelihood loss function as Basenji2^2^. We modified the dataset by extending the input sequence to 196,608 bp from the original 131,072 bp using the hg38 reference genome. The dataset contains 34,021 training, 2,213 validation, and 1,937 test sequences for the human genome, and 29,295 training, 2,209 validation, and 2,017 test sequences for the mouse genome. For the human genome, each example contains 2,131 transcription factor (TF) ChIP-seq, 1,860 histone modification ChIP-seq, 684 DNase-seq or ATAC-seq, and 638 CAGE tracks (total 5,313, **Supplementary Table 2**). For the mouse genome, each example contains 308 TF ChIP-seq, 750 histone modification ChIP-seq, 228 DNase-seq or ATAC-seq, and 357 CAGE tracks (total 1,643, **Supplementary Table 3**).

To train a model simultaneously on human and mouse genomes, we alternated between a batch containing data from the human genome and the mouse genome. The main Enformer model with 1,536 channels was trained on 64 TPU v3 cores with batch size of 64 (1 per core) using all-reduce gradient aggregation across the cores at every step. Batch normalization statistics were also aggregated across multiple replicas using 0.9 momentum. We used the Adam optimizer from Sonnet v2^40^ with a learning rate of 0.0005 and default settings for other hyperparameters: beta1=0.9, beta2=0.999, epsilon=1e-08. The optimal learning rate was discovered by grid search yielding the highest performance on the validation set. We linearly increased the learning rate from 0 to target value in the first 5,000 steps of training. We clipped gradients to a maximum global norm of 0.2. We used the same data augmentation as Basenji2^2^ during training by randomly shifting the input sequence by up to 3 bp and reverse-complementing the input sequence while reversing the targets. Finally, we fine-tuned the Enformer model on human data for 30k steps using a lower learning rate of 0.0001.

For the ablation studies and hyperparameter sweeps shown in **Extended Data Fig. 2**, we used 768 channels (1/2 of the original Enformer model obtained by using value size of 96 in MHA), a shorter 131 kb input sequence, and batch size of 32 trained on 32 TPU v3 cores. We did not fine-tune these models on the human data. For models using dilated convolutions instead of transformer blocks, we used a higher learning rate of 0.02 without learning rate ramp up. As for Enformer, the optimal learning rate was discovered by grid search yielding the highest performance on the validation set. All models were trained for 500k steps while only storing the model with the highest Spearman correlation of CAGE TSS gene expression across genes averaged across experiments computed on the validation set every 1,000 steps.

We used the validation set for hyperparameter selection and the test set for Basenji2 comparison (performed once). We considered two evaluation metrics: 1) Pearson correlation computed across all 128 bp binned genomic positions in the validation/test set for each output track. 2) Pearson correlation of CAGE gene expression values (log(1+x) transformed and standardized across genes for each experiment) of all protein-coding genes in the validation/test set computed either for each CAGE experiment across genes (main metric) or across CAGE experiments for each gene (shown in **Fig. 1b**). Observed and predicted gene expression values were obtained by summing up the observed/predicted CAGE read counts at all unique TSS locations of the gene. For each TSS location, we used the 128 bp bin overlapping the TSS as well as the two neighbouring bins (i.e. 3 bins in total). We used test-time augmentation during model evaluation: we averaged the predictions from 8 sequences augmented the same way as during training (≤3 bp shifts and reverse-complementation). We only evaluated the performance of our model on the test set once to generate **Fig. 1** and have not used the test set during model development.

### Experimental-level accuracy

To estimate the experimental-level accuracy of gene expression measurements, we used 204/638 CAGE samples that had multiple replicate experiments (589 experiments in total). For each CAGE sample, we divided the replicate experiments into two groups such that the total number of reads was roughly equal in each group (**Supplementary Table 1**). For each group, we summed the reads for all replicates to generate a pseudo-replicate. Replicate level accuracy was computed by treating data from pseudo-replicate 1 as predictions and data from pseudo-replicate 2 as targets.

Since only 204/638 CAGE samples had multiple replicates, we developed a computational approach to estimate the replicate level accuracy of all 638 CAGE samples. Since many CAGE tracks are highly correlated, a linear combination of other tracks can be used to estimate the replicate experiment of a particular CAGE track. For each track *t*, we trained a linear model on the same training set as the Enformer model using batch-normalized input values of all other tracks t’≠t and their log(1+x) transformed version, followed by softplus activation. Performance of this model was evaluated on the same test set as the Enformer model.

### Enhancer prioritization

We obtained a set of enhancer–gene pairs tested using a CRISPRi assay perturbing the enhancer of interest while measuring the expression change of the gene in K562 cells from two studies: Gasperini et al 2019^8^ using scRNA-seq to measure expression changes, and Fulco et al 2019^11^ using Flow-FISH. We transformed the enhancer and gene coordinates from hg19 to hg38 using the UCSC liftOver web tool^41^. Each enhancer–gene pair contains a label denoting whether or not a significant expression change was induced after CRISPRi treatment. We denoted the set of all enhancers as ‘candidate’ enhancers and those that showed a change in expression as ‘validated’ enhancers. We evaluated different methods on their ability to classify or prioritize enhancer–gene pairs that exhibited a significant expression change using area under Precision-Recall Curve (auPRC)^11^.

To prioritize enhancer–gene pairs with sequence-based models, we computed three different scores: gradient*input, attention, and *in silico* mutagenesis (ISM). For each enhancer–gene pair, we determined the major TSS of the gene by taking the highest predicted CAGE value in K562 using Enformer. We extracted the DNA sequence centered at the main TSS and computed the following different enhancer-gene scores:

- **gradient*input**: We computed the absolute value of the gradient of the CAGE targets (either using the K562-specific CAGE targets or all CAGE targets, **Extended Data Fig. 6c**) at the TSS w.r.t the input sequence. At each sequence position, the gradient of the observed base was used. Note that since our input sequence is one-hot encoded, taking the input gradient of the non-zero channel (the input sequence base), is equivalent to computing gradient*input attributions^12^. We note that “CAGE at TSS” always means summing the absolute gradient values from 3 adjacent bins, as is also done in gene-focused model evaluation. The 3 bins include the bin overlapping the TSS and one flanking bin on each side. Enhancer–gene score was obtained by summing the absolute gradient*input scores in the 2 kb window centered at the enhancer.
- **attention**: We first averaged transformer attention matrices across all heads and layers. We then extracted the row corresponding to the query index positioned at TSS with keys corresponding to different spatial positions, denoting how much the model attended to these positions when making the predictions at the TSS. We only computed this contribution score for Enformer. Enhancer–gene score was obtained by summing the attention scores in the 2 kb window centered at the enhancer.
- **ISM:** The *in silico* mutagenesis enhancer-gene score was computed by comparing K562 CAGE predictions at the TSS from the reference sequence with predictions from modified sequence where the 2 kb enhancer sequence was replaced by a random sequence: |f(modified) - f(reference)|.

To reproduce the ABC score introduced in Fulco et al^11^, we obtained the BigWig of H3K27ac ChIP-seq data in K562 from ENCODE with file accession ENCFF779QTH and DNase with file accessions ENCFF413AHU and ENCFF936BDN (we summed the normalized reads from these two replicates). For each track and enhancer, we summed up the signal at the enhancer in a fixed window of 2 kb centered at the enhancer. This fixed and broader window yielded better performance compared to the variable window size of ∼500 bp as used in the original ABC score (**Extended Data. Fig. 4a**).

### TAD boundary attention

We obtained a list of topologically associating domain (TAD) boundaries measured in IMR90 cells following the analysis of Krietenstein et al^42^. We randomly selected 1,500 TAD boundaries, extracted the reference genome sequence centered at these positions, made predictions with Enformer, and obtained the transformer attention matrices. We averaged the attention matrices across all sequences, layers, and transformer heads (yielding a 1,536×1,536 matrix). We did the same for 1,500 sequences from our validation set; these sequences do not contain any specific alignment to a particular regulatory element and can hence be considered as ‘background’. We then computed the difference between the two attention matrices (attention_tad_boundary - attention_other) and visualized it as a heatmap. “Query” denotes the position from which we are attending to all other positions in the sequence called “keys”. To compute the statistical significance of the patterns observed in the heatmap, we incrementally partitioned the attention matrix into the following segments:

- Query at insulator: |key| < 6 kb (6kb corresponds to 48 bins of 128 bp)
- Key at insulator: |query| < 6 kb
- Diagonal: |query - key| < 6 kb
- Within TAD: sign(query*key) = 1
- Across TAD: sign(query*key) = -1

Each segment also excludes segments from previous segments thereby making sure that none of the segments overlap (for example “Diagonal” segment will not contain regions overlapping “Key at insulator” and “Query at insulator”). For the attention matrix of each sequence (averaged across layers and heads), we computed the average attention value in the segment of interest.

### GTEx SLDP

We predicted the effect of a genetic variant on various annotations by computing a forward pass through the model using the reference and alternative alleles, subtracting their difference, and summing outputs across the sequence to obtain a signed score for each training dataset. We averaged scores computed using the forward and reverse complement sequence and small sequence shifts to the left and right. We computed scores for all 1000 Genomes SNPs.

Signed linkage disequilibrium profile (SLDP) regression is a technique for measuring the statistical concordance between a signed variant annotation *v* and a genome-wide association study’s marginal correlations 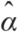 between variants and a phenotype^22^. The functional correlation between *v* and the true variant effects on the phenotype describes how relevant the annotation is for the phenotype’s heritability. Our model produces these signed variant annotations. SLDP estimates this functional correlation using a generalized least-squares regression, accounting for the population linkage disequilibrium structure. It performs a statistical test for significance by randomly flipping the signs of entries in *v* in large consecutive blocks to obtain a null distribution. We follow previous work in conditioning on background annotations describing minor allele frequency and binary variables for variant overlap with coding sequence (and 500 bp extension), 5′ UTR (and 500 bp extension), 3′ UTR (and 500 bp extension), and introns.

We used GTEx v7a summary statistics for 48 tissues that had been preprocessed for SLDP^21^. In these statistics, each SNP’s effect is summarized over all cis-genes using the following transformation 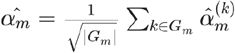 where *G*_*m*_ is the set of all genes for which a cis-eQTL test was performed for variant *m* and is the marginal correlation of SNP *m* and 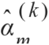 gene *k* expression^22^. We passed 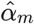 to SLDP for analysis of variant predictions.

### Fine-mapped GTEx classification

To study specific eQTLs without need for considering LD, we performed statistical fine-mapping of GTEx v8 using the SuSiE method^21,24^. We focused on variants with posterior inclusion probability (PIP) in a credible causal set >0.9, which ranged from a minimum of 166 variants for substantia nigra to 2,740 for tibial nerve. We arranged a classification task to discriminate between these positive causal variants and a matched set of negative variants. When available, we chose a negative variant matched to each causal variant from the set with PIP<0.01 but |z-score|>4 tested for the same gene. When unavailable for the same gene, we chose from the set with PIP<0.01 and |z-score|>6 genome-wide.

To determine how informative different variant annotations are, we trained separate random forest classifiers for each tissue to distinguish causal from non-causal variants using eight-fold cross-validation. We selected the default hyperparameters of the scikit-learn 0.22 implementation after finding negligible accuracy gains from modifying them^43^. However, due to the large number of features derived from the training datasets, setting the maximum features considered per decision tree split to log_2_ of the total number of features greatly improved the computational efficiency. We fit 100 iterations of stochastic cross-validation shuffling and random forest fitting to delineate a low-variance estimate of model accuracy. We performed statistical tests comparing two different model feature sets by comparing the 8*100 distinct test set auROCs.

For signed GTEx analysis, we benchmarked model predictions based on their ability to discriminate causal variants that increase versus decrease gene expression. In this analysis, we removed variants that affect gene expression in opposite directions for different cis-genes. We manually matched FANTOM5 CAGE sample descriptions to the GTEx tissues. We skipped cases with more than three possible matches. In cases with two or three possible matches, we chose the CAGE sample with the best average concordance between the Basenji2 and Enformer predictions. We computed auROC statistics by ranking causal variants by their signed prediction for that sample.

### Dimensionality reduction of variant effect scores

We used principal component analysis (PCA) to reduce 5,313 variant effect features from Enformer to 20 principle components. We used variant effect scores from 1000 Genomes SNPs on chromosome 9 and performed the following steps: 1. subtracted the median and divided by standard deviation estimated from the interquartile range as implemented in RobustScaler in scikit-learn (v0.23.2), 2. reduced the dimensionality to 20 principle components using TruncatedSVD from scikit-learn, and 3. normalized the resulting principal component features using RobustScaler to obtain z-scores.

### Benchmarking variant effect predictions on saturation mutagenesis data

We acquired training and test sets as well as the predictive accuracies of individual competition participants from the CAGI5 competition^26^ (M. Kircher, personal communication, https://genomeinterpretation.org/content/expression-variants). For each variant and locus, we evaluated its effect as the predicted difference between the reference and alternative allele summed in four flanking bins representing 512 bp, producing 5,313 features in the human. All CAGE features were log-transformed after adding a pseudocount of 1 prior to computing this difference. For each allele, we averaged predictions for the forward and reverse-complemented sequence. We scaled the features from the test set with scaling factors computed on the features from the training set, such that the training features had a mean of 0 and standard deviation of 1. Following our previous work^44^, we then trained a lasso regression model for each locus using these features and the corresponding training set. The strength of the regularization was controlled by a single λ parameter, which was optimized using 10-fold cross-validation for each training set using the *cv*.*glmnet* function of the *glmnet* library in R.

For our training-free comparisons, we selected the subset of features corresponding to cell-type matched and cell-type agnostic predictions of changes in CAGE and DNase. For the cell-type agnostic models, we used the subset of all 638 CAGE or 674 DNase features (**Supplementary Table 2**). For the cell-type matched models, we additionally required the CAGE/DNase features to contain the following substrings: i) “HepG2” for F9, LDLR, and SORT1, ii) “K562” for GP1BB, HBB, HBG1, and PKLR, iii) “HEK293” for HNF4A, MSMB, TERT (performed in HEK293T cells), and MYCrs6983267. For several loci, a perfectly matched DNase or CAGE sample did not exist. We therefore selected the most closely matched feature based on the following substrings: i) “pancrea” for ZFAND3, ii) “glioblastoma” for TERT (performed in GBM cells), iii) “eratinocyte” for IRF6, and iv) “SK-MEL” for IRF4. For each locus, we extracted the features matching the aforementioned substrings, and used the first principal component (PC) of the indicated features as our summary statistic, inverting the sign of the PC if it was negatively correlated to the mean of the features.

## Supplemental figure

**Extended Data Fig. 1:**
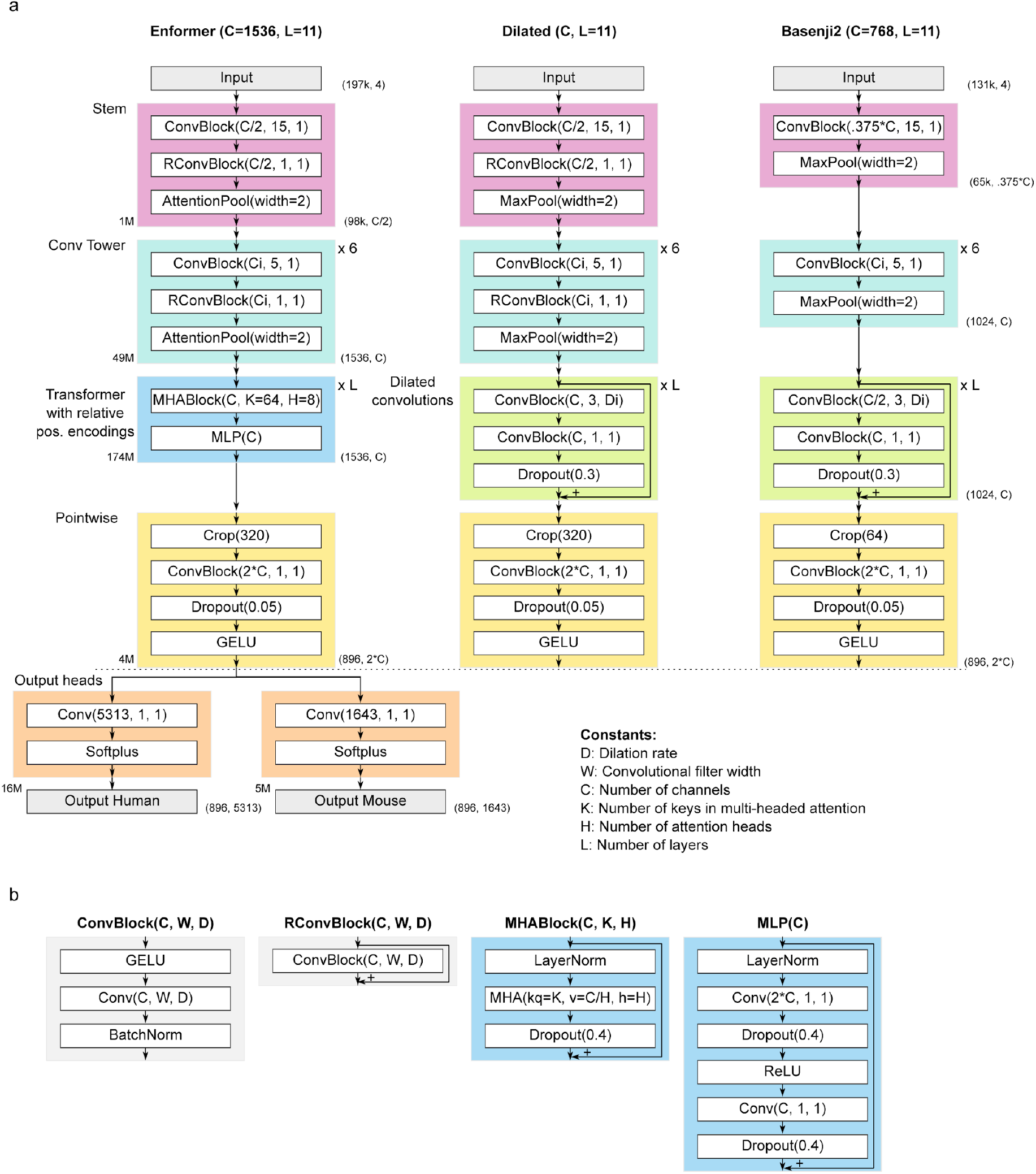
Enformer model architecture and comparison to Basenji2. **a)** From left to right: Enformer model architecture, ‘dilated’ architecture used in ablation studies obtained by replacing the transformer part of the model with dilated convolutions, and Basenji2^2^. Output shapes (without batch dimensions) are shown as tuples on the right side of the blocks. The number of trainable parameters for different parts of Enformer are shown on the left side of the blocks. The two main hyperparameters of the model are the number of transformer/dilated layers, L, and the number of channels, C. All models have the same two output heads as shown on the Enformer at the bottom. The number of channels in the convolutional tower Ci was increased by a constant multiplication factor to reach C channels starting from C/2 (or 0.375*C for Basenji2) in 6 layers. For dilated layers, we increased the dilation rate Di by a factor of 1.5 at every layer (rounded to the nearest integer). **b)** Definition of different network blocks in terms of basic neural network layers. MHA denotes multi-headed attention using relative positional encodings with *kq* representing the number of key/query size, *v* representing the value size and *h* the number of heads. Number of relative positional basis functions is equal to value size *v*.

**Extended Data Fig. 2:**
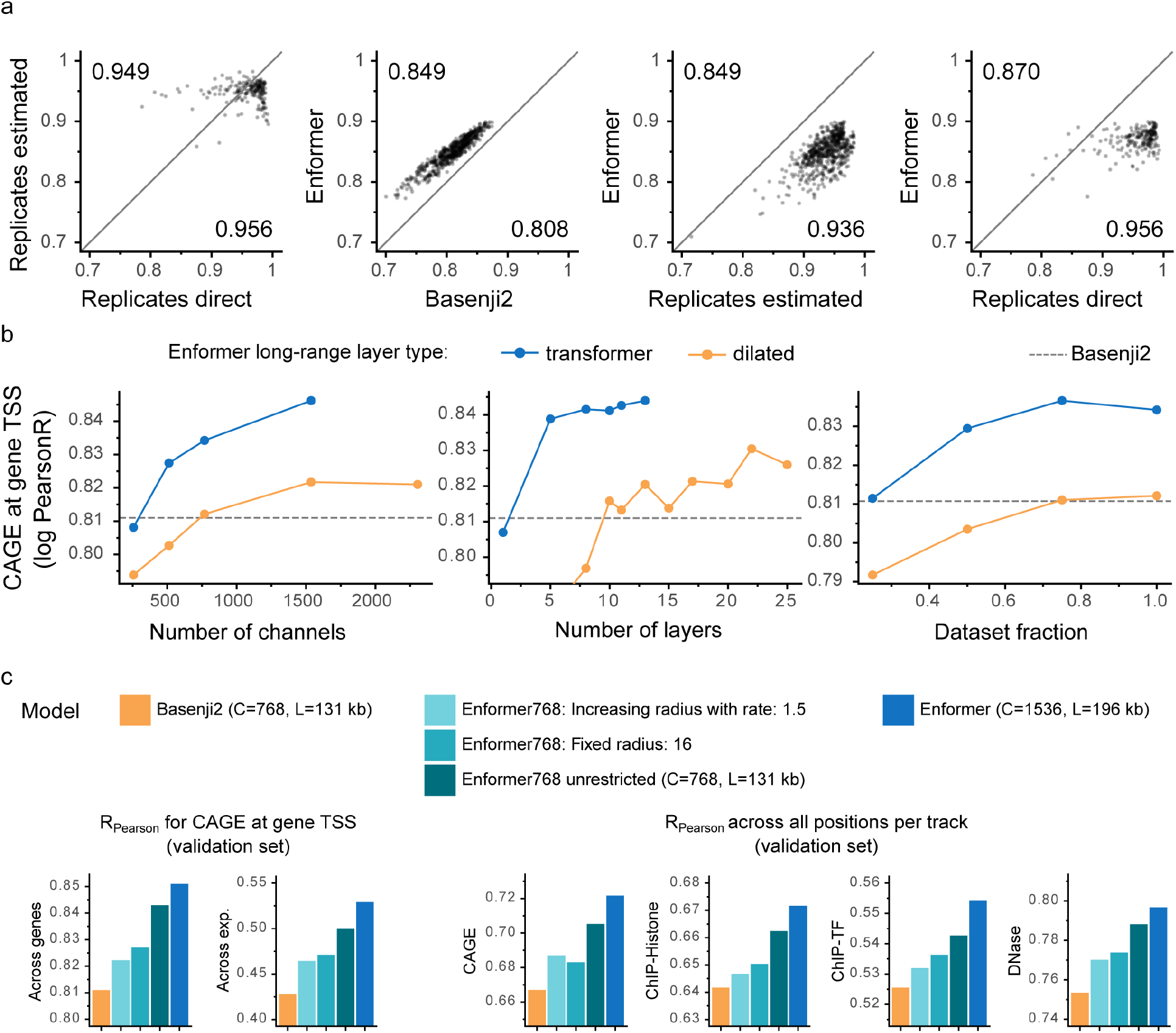
Replicate level accuracy, comparison to dilated convolutions. **a)** Gene expression correlation (log(1+x) pearsonR) for each CAGE track across protein-coding genes comparing experimental-level accuracy computed in two ways (estimated and direct) to Enformer. For “Replicates direct”, CAGE replicate experiments were partitioned into two groups and compared against each other. For “Replicates estimated”, a predictive model was used to impute CAGE values of a particular track from all other tracks. **b)** Enformer with original transformer layers (**Extended Data Fig. 1a** left) performs better than Enformer with dilated convolutions (**Extended Data Fig. 1a** center) across different model sizes and training dataset subsets as measured CAGE gene expression correlation in the validation set (same metric as in **Fig. 1b** across genes). At 15 dilated layers, the model starts to reach outside of the input sequence range (receptive field of 224,263 bp). Note that all the evaluations here are limited by TPU memory preventing you from using more layers or channels. **c)** Performance comparison to Basenji2 (left) and Enformer (right) to Enformer with the same receptive field (44 kb) as Basenji2 by either allowing a fixed attention radius of 16 across all layers where query can attend to at most 16 positions away (Enformer 769: Fixed radius: 16) or by exponentially increasing the respective field in the same way as the dilation rate in Basenji2. Enformer768 was trained with the same number of 768 channels and 131 kb input sequences as Basenji2, whereas Enformer uses two times more channels and 1.5 times longer sequence. Same evaluation metrics are shown as in **Fig. 1**.

**Extended Data Fig. 3:**
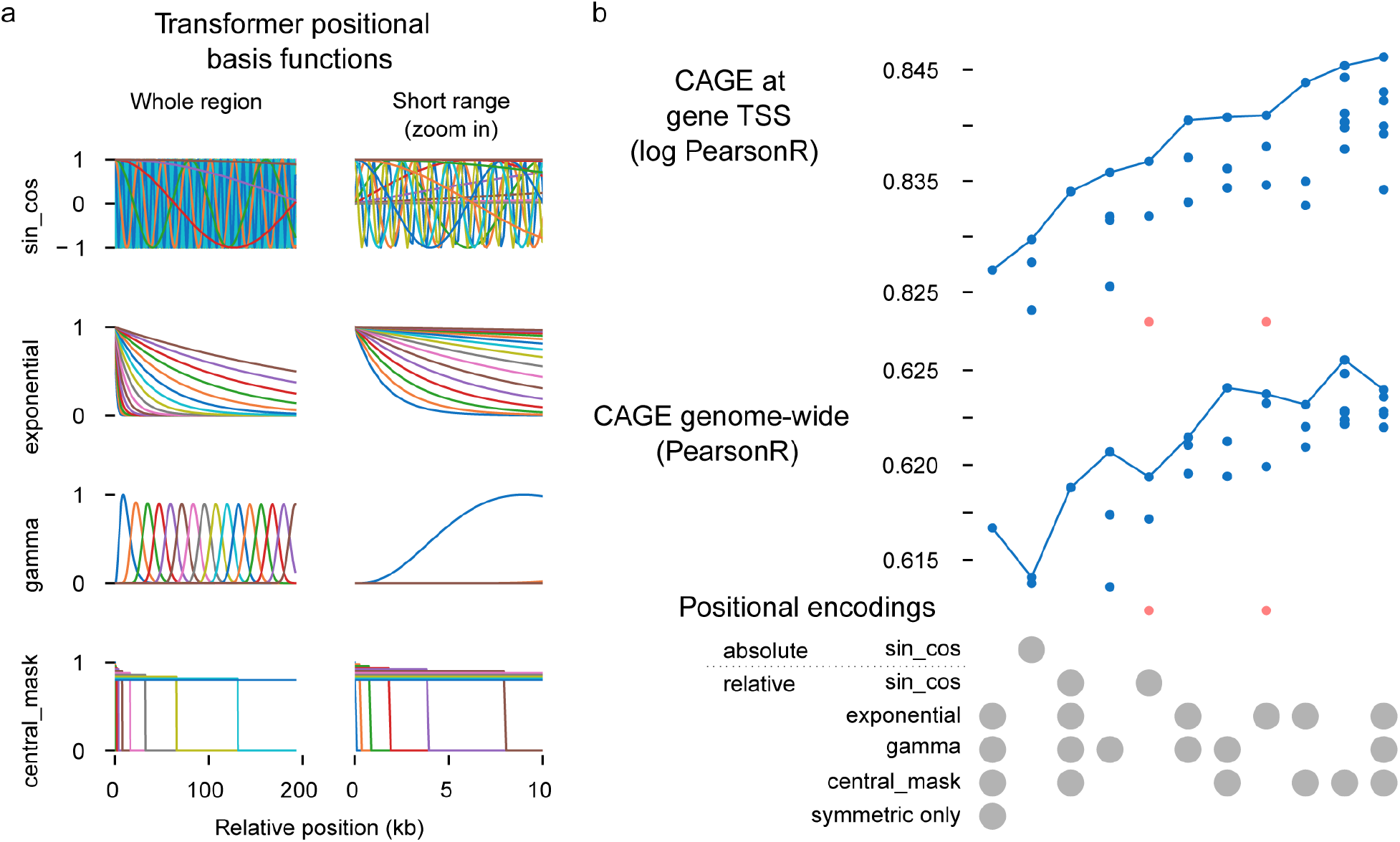
Custom relative positional encoding functions are required for good predictive performance. **a)** Relative positional encoding basis function options for the transformer model. Sine/cosine basis functions are frequently used in the NLP literature for both absolute or relative positional encodings. Enformer uses a concatenation of exponential, gamma and central_mask relative positional encodings. For each basis function, a symmetric f(|x|) and asymmetric sign(x) * f(|x|) basis function will be used to introduce directionality and thereby inform the model of what is upstream or downstream of the TSS. **b)** Validation set performance as measured by CAGE gene expression correlation across protein coding genes (top; same metric as in **Fig. 1b** across genes) or across all positions (bottom; same metric as displayed in **Fig. 1c** CAGE) for models trained with different classes of positional encoding functions in the transformer. Custom relative positional encodings show better performance than using standard sin/cos basis functions or using absolute positional encodings, likely because they can better capture the decreasing importance of enhancers with increased distance. Also, symmetric only (f(|x|) version shows much lower performance than using both, symmetric and asymmetric versions. All models use the same 96 total number of basis functions. Each positional encoding configuration was trained with multiple different random seeds. Red points denote runs with lower performance than the y-axis limits.

**Extended Data Fig. 4:**
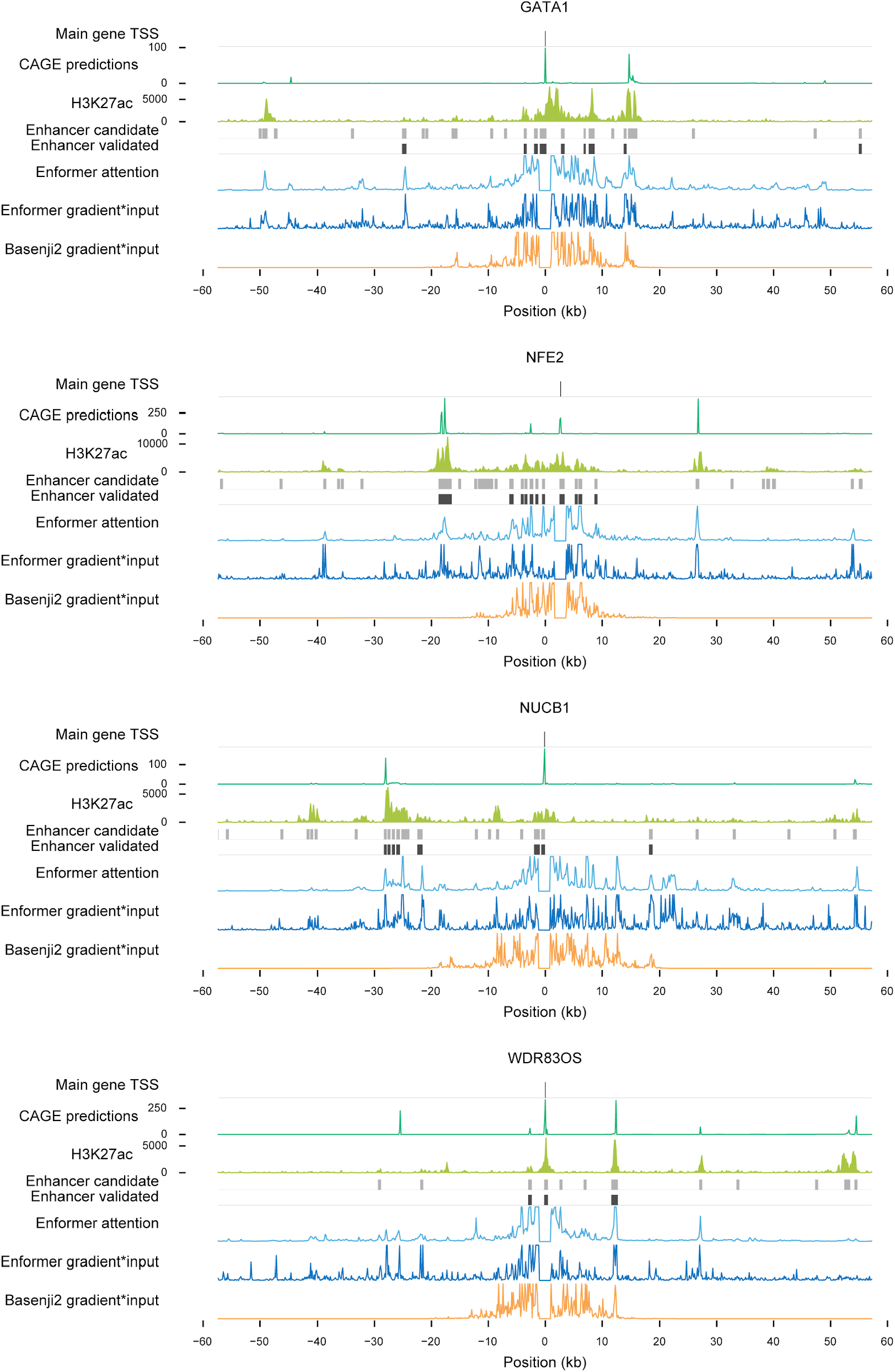
Example loci from Fulco *et al* 2019. Other example loci from Fulco et al 2019 as shown in Fig. 2.

**Extended Data Fig. 5:**
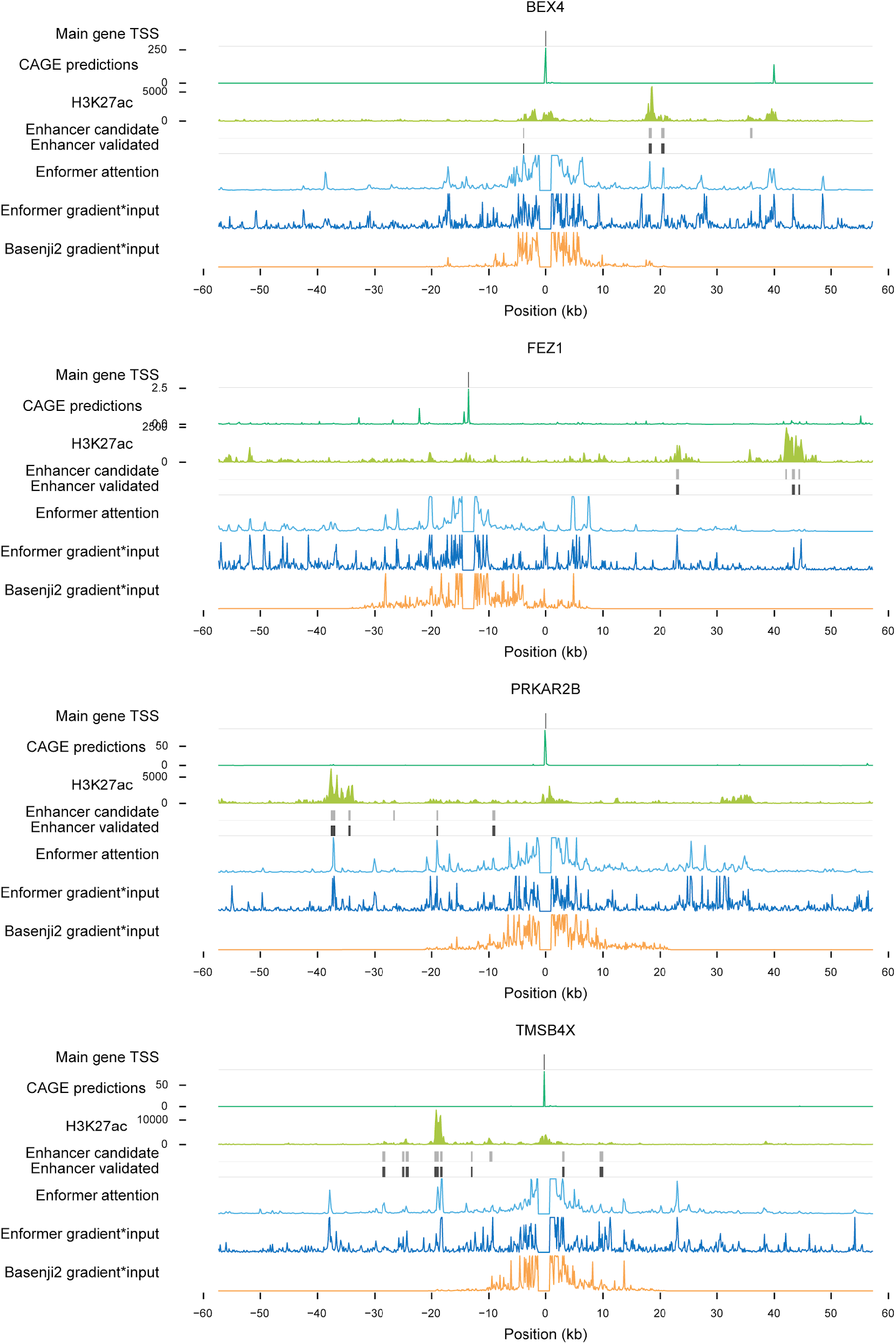
Example loci from Gasperini *et al* 2019. Other example loci from Gasperini et al 2019 as shown in Fig. 2.

**Extended Data Fig. 6:**
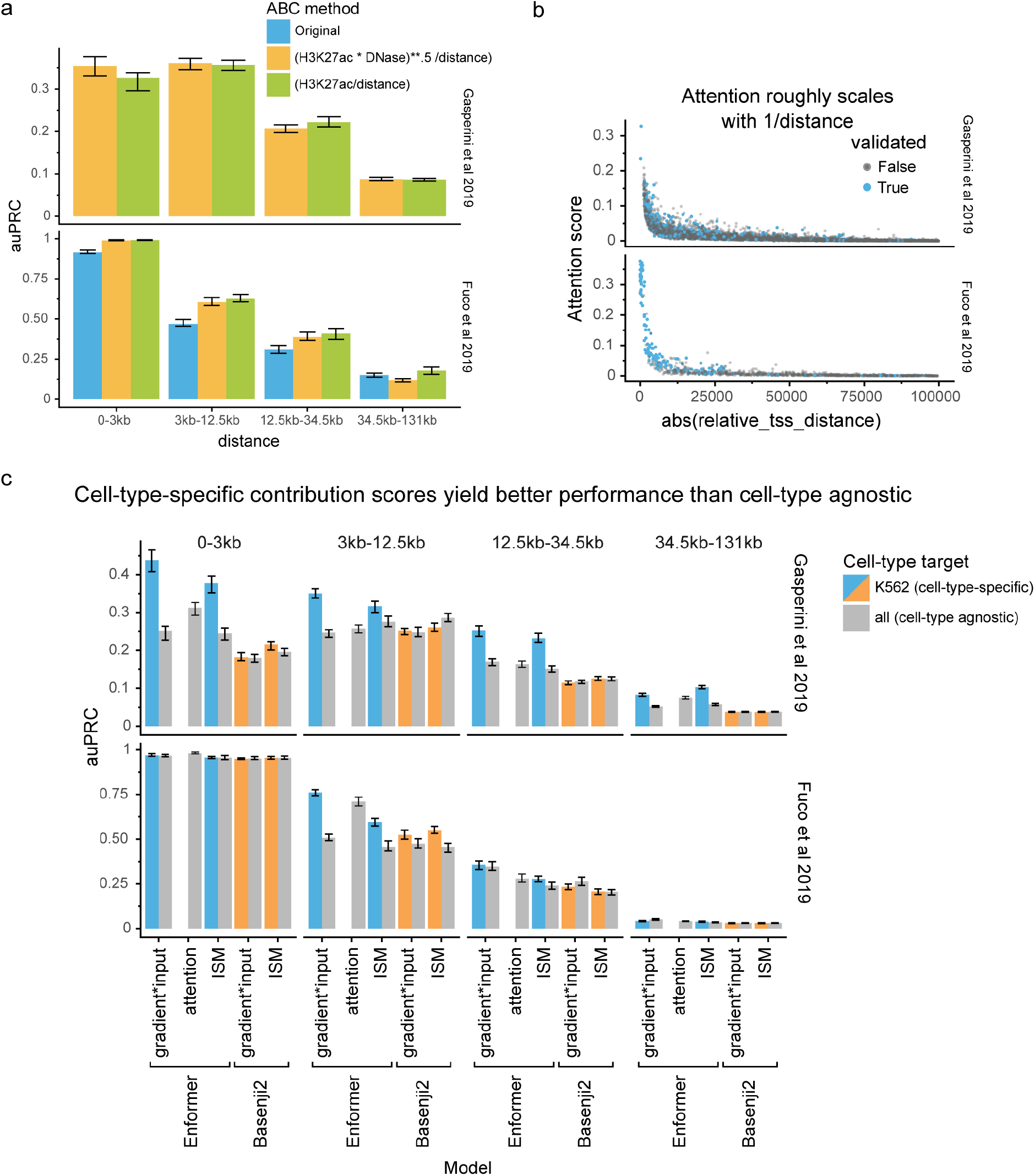
Tissue-specific contribution scores are required for good enhancer-recall performance. **a)** Enhancer–gene ranking performance comparison of different ABC score versions, including the original score which uses DNase, H3K27ac, and Hi-C data. H3K27ac / distance is a good if not even slightly better proxy for the ABC score. The alternative versions use a fixed and 2 kb wide aggregation window whereas the original uses a dynamic peak width depending on the DNase peak width. **b)** Attention-based contribution score as a function of distance at all enhancer–gene pairs in both studied datasets. **c)** Contribution scores that are cell-type-specific (shown in green, achieved by computing the contribution scores w.r.t. cell-type-specific target variables) outperform cell-type agnostic contribution scores (shown in orange).

**Extended Data Fig. 7:**
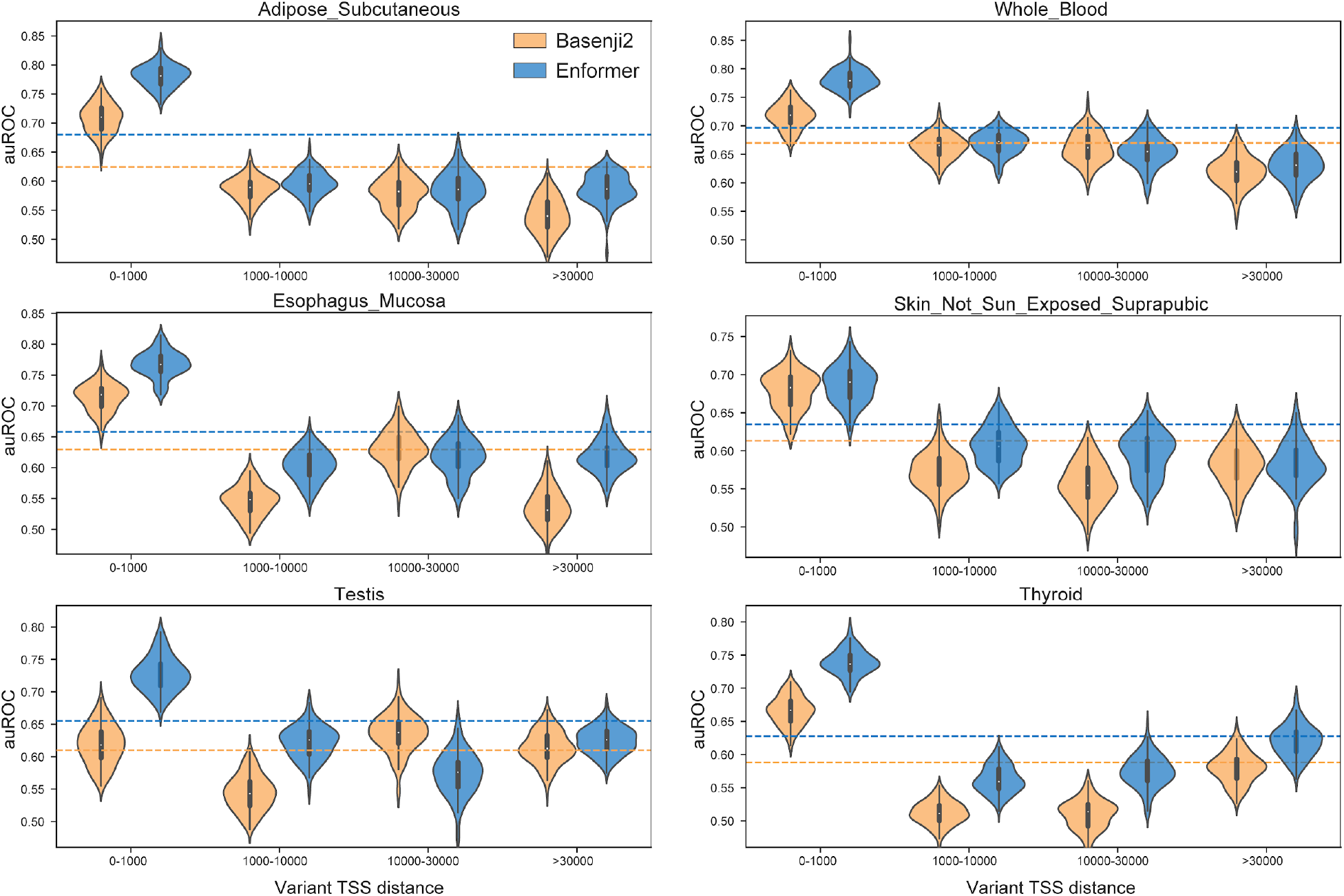
Enformer outperforms Basenji2 on eQTL sign prediction. For each of the GTEx tissues, we manually matched FANTOM5 CAGE sample descriptions to choose a single matched dataset (Methods). We then arranged a classification task to discriminate between fine-mapped causal eQTLs in which the minor allele increases gene expression versus eQTLs in which the minor allele decreases gene expression. We computed auROC statistics by ranking causal variants by their signed prediction for the corresponding sample. To consider the influence of variant distance to TSS, we compute auROC in four bins of roughly equal size. Across tissues and TSS distances, Enformer predictions usually achieve more accurate classification of eQTL sign than Basenji2 predictions. We display six example tissues with large numbers of fine-mapped eQTLs and with clear correspondence between CAGE and GTEx tissues. Violin plots show the auROC distribution of 100 bootstrap samples from the full set of variants. Both models struggle with variants beyond the promoter, highlighting an important problem for future research.

## Supplementary Tables

**Supplementary Table 1:** Individual CAGE replicate experiments. Columns ‘replicate_group’ denotes the CAGE sample and ‘rep_id’ denotes the individual replicates. Only replicate groups with multiple replicates (multiple_reps=True) were used to compute the experimental-level accuracy. Division of individual replicates for each sample (replicate_group) into pseudo-replicates is denoted in the pseudo-rep columns.

**Supplementary Table 2**,**3:** Description of 5,313 human and 1,643 mouse tracks used to train and evaluate the model. Columns after assay_subtype mark which tracks were included for the cell-type or cell-line specific analysis in Fig. 4 (Methods).

